# Effects of Attentional Bias Modification on Residual Symptoms in depression. A Randomized Controlled Trial

**DOI:** 10.1101/318279

**Authors:** Rune Jonassen, Catherine J. Harmer, Eva Hilland, Luigi A. Maglanoc, Brage Kraft, Michael Browning, Tore C. Stiles, Vegard Ø. Haaland, Torkil Berge, Nils Inge Landrø

**Author notes:** **Correspondence**: Rune Jonassen, Clinical Neuroscience Research Group, Department of Psychology, Forskningsveien 3A, 0373 Oslo & Nils Inge Landrø, Clinical Neuroscience Research Group, Department of Psychology, Oslo, Norway. Diakonhjemmet Hospital, Division of Psychiatry, Oslo, Norway, Forskningsveien 3A, 0373, Oslo.

## Abstract

**Background:** Following treatment, many depressed patients have significant residual symptoms. However, large randomised controlled trials (RCT) in this population are lacking. When Attention bias modification training (ABM) leads to more positive emotional biases, associated changes in clinical symptoms have been reported. A broader and more transparent picture of the true advantage of ABM based on larger and more stringent clinical trials have been requested.

**Aims:** To evaluate the early effect of two weeks ABM training on blinded clinician-rated and self-reported residual symptoms, and whether changes towards more positive attentional biases (AB) would be associated with symptom reduction.

**Method:** A total of 321 patients with a history of depression were included in a preregistered randomized controlled double-blinded trial. Patients were randomised to an emotional ABM paradigm over fourteen days or a closely matched control condition. Symptoms based on the Hamilton Rating Scale for Depression (HRSD) and Beck Depression Inventory II (BDI-II) were obtained at baseline and after ABM training.

**Results:** ABM training led to significantly greater decrease in clinician-rated symptoms of depression as compared to the control condition. No differences between ABM and placebo were found for self-reported symptoms. ABM induced a change of AB towards relatively more positive stimuli associated with greater symptom reduction.

**Conclusion:** The current study demonstrates that ABM produces early changes in both AB and blinded clinician-rated depressive symptoms. ABM may have practical potential in the treatment of residual depression.

*ClinicalTrials.gov ID: NCT02658682*

## BACKGROUND

A wide range of treatments are available for depression. However, systematic reviews point to limited efficacy in terms of remission, response rates and long-term effects for both pharmacological (1) and psychological treatments (2). Forty percent of patients do not, or only partially, respond to treatment, and less than one third are completely recovered after treatment (3). Residual symptoms following treatment are associated with decreased return to function, reduced quality of life and increased risk of recurrence highlighting the need for further intervention (4).

Depression is associated with an increased focus on negative interpretations of events and negative biases in attention and memory. Previous studies have reported that clinically depressed subjects orient their attention toward negative faces rather than neutral or positive faces (5). Biases towards negative faces have also been reported in previously depressed, currently euthymic subjects (6), and in never-depressed individuals with a family history of depression (7). In a seminal experimental study a causal role for negative attention biases in the expression of depressive and anxious symptoms was demonstrated (8). They found higher levels of anxiety and depression related mood ratings in undergraduates, as a reaction to a stressor, after being trained to preferentially attend to negative versus neutral stimuli. Together these results indicate that negative attentional biases (AB) may constitute causative, vulnerability and maintenance factors rather than simple markers of lowered mood. This suggests that interventions designed to reduce AB may act to reduce symptoms in patients treated for depression.

Computerized Attention Bias Modification (ABM) procedures aim to shift emotional biases towards more positive and less negative stimuli. ABM is low on resource requirements and easy to disseminate and might therefore be widely used by patients with residual symptoms or while waiting for more resource intensive interventions like psychotherapy. While some studies have reported an effect of ABM in depression a number of meta-analyses have suggested a small effect size, although definitive conclusions have been limited by small sample sizes and poor trial methodology employed in many studies (9-11). In addition, some potential techniques for altering AB have failed to modify AB as part of the treatment programme, which limits their ability to decrease depressive symptoms (12), but see also (13).

The main objective of our study was to test the early efficacy of ABM in a large group of previously depressed patients, but with residual symptoms using a preregistered trial design. We hypothesized that two weeks of ABM training would reduce clinician- and self-reported residual symptoms of depression, as compared to a matched control condition. Given the clear mechanistic role for altered bias in the clinical effects of ABM, we predicted a change towards more positive biases in the ABM group and that this change would be associated with decreased symptoms.

## METHODS

### Study design and Procedure

Patients who had previously been identified as suffering from depression were randomised to receive two weeks of either ABM- or placebo training. Participants completed two sessions of ABM daily using laptops provided by the research team. A detailed calendar was used to specify the scheduling of the training sessions for each participant. Symptoms of depression and AB were assessed at baseline and immediately after the intervention.

### Participants

The main recruitment base was an outpatient clinic in the Department of Psychiatry, Diakonhjemmet Hospital in Oslo. Participants were also recruited from other clinical sites, by local advertisements, and via social media. Candidates were pre-screened by phone for exclusion criteria before in person formal clinical evaluation and enrollment. Individuals diagnosed with current- or former neurological disorders, psychosis, bipolar spectrum disorders, substance use disorders, attention deficit disorder, and head trauma were excluded.

A total of 377 participants between 18-65 years old were recruited for clinical evaluation. Participants had experienced more than one previous episode of depression as defined using the Mini International Neuropsychiatric Interview, version 6.0.0. (MINI). A total of 56 subjects were excluded following the clinical evaluation. The period of recruitment and follow-up was January 2015 to October 2016 and was first posted in *ClinicalTrials.gov* in January 2016, and was therefore partly retrospectively registered.

A total of 321 participants with a history of Major Depressive Episodes (MDE) were randomised to receive ABM or placebo. Thirty-seven participants who fulfilled the MINI criteria for current MDE were enrolled onto the study in error (current MDE was described as exclusion criteria in the pre-registered protocol). Inclusion of data from these participants did not influence the results (see supplementary analysis).

### Attention Bias Modification Procedure

The ABM task was a computerized visual dot-probe procedure adopted from (14). Paired images of faces (the stimuli), were presented followed by one or two dots (a probe), which appeared behind one of the stimuli. Participants were required to press one of two buttons as quickly as possible to indicate the number of dots in the probe. The types of stimuli used during the ABM task were pictures of emotional faces of three valences; positive (happy), neutral, or negative (angry and fearful). A single session of the task involved 96 trials with equal numbers of the three stimulus pair types. In addition, there were equal numbers of trials in which the stimuli were randomly presented for 500- or 1000 ms before the probe was displayed. In each trial of the task, stimuli from two valences were displayed, in one of the following pairing types: positive-neutral, positive-negative, and negative-neutral. In the ABM condition, probes were located behind positive (valid trials) stimuli in 87 % of the trials, as opposed to 13% with probes located behind the more negative stimuli (invalid trials). Thus, when completing the ABM, participants should learn to deploy their attention toward the relatively more positive stimuli, and in this way develop a more positive AB. The neutral ABM control (placebo) condition was identical in every respect, other than the location of the probe, which was located behind the positive (valid trials) stimuli in only 50% of the trials. Participants completed two sessions of ABM daily during the course of fourteen days (28 sessions in total) on identical notebook computers that were set up and used exclusively for ABM-training.

### Measurement of Attentional Bias

AB was measured at baseline and after two weeks of ABM using a standard visual probe procedure consisting of 96 trials, with the same trial types as used in the ABM procedure. Novel facial stimuli were used in the assessment tasks. AB was calculated as the difference in reaction between trials in which the probe replaced the relatively more negative face vs. the more positive face ([(SUM(positive up-probe down, positive down, probe up) - SUM(positive up-probe up, positive down-probe down)]/2.). Thus a more positive score reflects a greater bias towards the more positive stimuli.

### Randomization and blinding

An independent lab technician (not involved in the day to day collection of data) prepared training computers to deliver either ABM or placebo treatment according to a randomization list in a 1:1 ratio ensuring that allocation was concealed from all researchers involved in screening procedures and all participants. Allocation to intervention is stored in the ABM raw data file that was opened and merged with screening data for statistical analyses first after the data collection period was finished in October 2016. Outcome assessors and participants were therefore blind to allocation during the whole study.

### Study outcome

The primary outcome measures were The Hamilton Rating Scale for Depression (HRSD) (15) and Beck Depression Inventory (BDI-II) (16) administered to assess both clinician - and self evaluations of symptoms. Secondary outcome was AB measured by a single session of the placebo-training task at baseline and after two weeks follow-up. Comorbid anxiety symptomatology was screened by the use of The Beck Anxiety Inventory (BAI-II) (17).

### Statistical analysis

All data were analysed using PASW 25.0 (IBM). The primary outcomes for the Intention to treat (ITT) sample was analysed by using repeated measures ANOVA with intervention (ABM versus placebo) as a fixed factor. Symptom at baseline and at two weeks follow-up (HRSD or BDI-II) was entered as within subject variable (time). To test the interaction between changes in symptoms and changes in AB, statistically significant symptom changes between ABM and placebo were followed up by repeated measures ANOVA by adding AB (two weeks follow-up minus baseline across valences and stimulus durations) as a second time factor in the model. Thereafter, the individual distribution of symptom change and AB was explored within groups with linear regression models.

A requirement of the national ethical committee was that study participants could elect to have all data stored on them during the study erased (i.e. including randomisation information) which precludes imputation of missing data. Of the 20 participants who did not complete the study following randomisation, 10 elected to have their data erased and thus data was imputed only for the remaining 10 remaining allocated to intervention. Thus, Thus the ITT analyses included 311 participants while participant with full data sets included 301 participants (see supplementary Figure 1.). A detailed description of procedures for data reduction, imputation and a post hoc analysis of the potential impact of noncompliance, stimulus valence-and duration, and sensitivity analysis are provided as supplemental information.

## RESULTS

### Sample characteristics

### Primary outcomes

There was a statistically significant difference (time) between ABM and the control condition in blinded clinician-rated symptoms as measured by the HRSD [F (1,309) = 6.78, η^2^ = .02, p < .01]. Means and standard deviations at baseline were 9.2 (5.9) for ABM and 8.3 (5.0) for placebo and changed to 8.3 (5.9) and 8.8 (5.7) at two weeks follow-up. Together the results show a positive change after ABM as compared to placebo at two weeks follow-up (Figure 1.).

**Figure 1.**
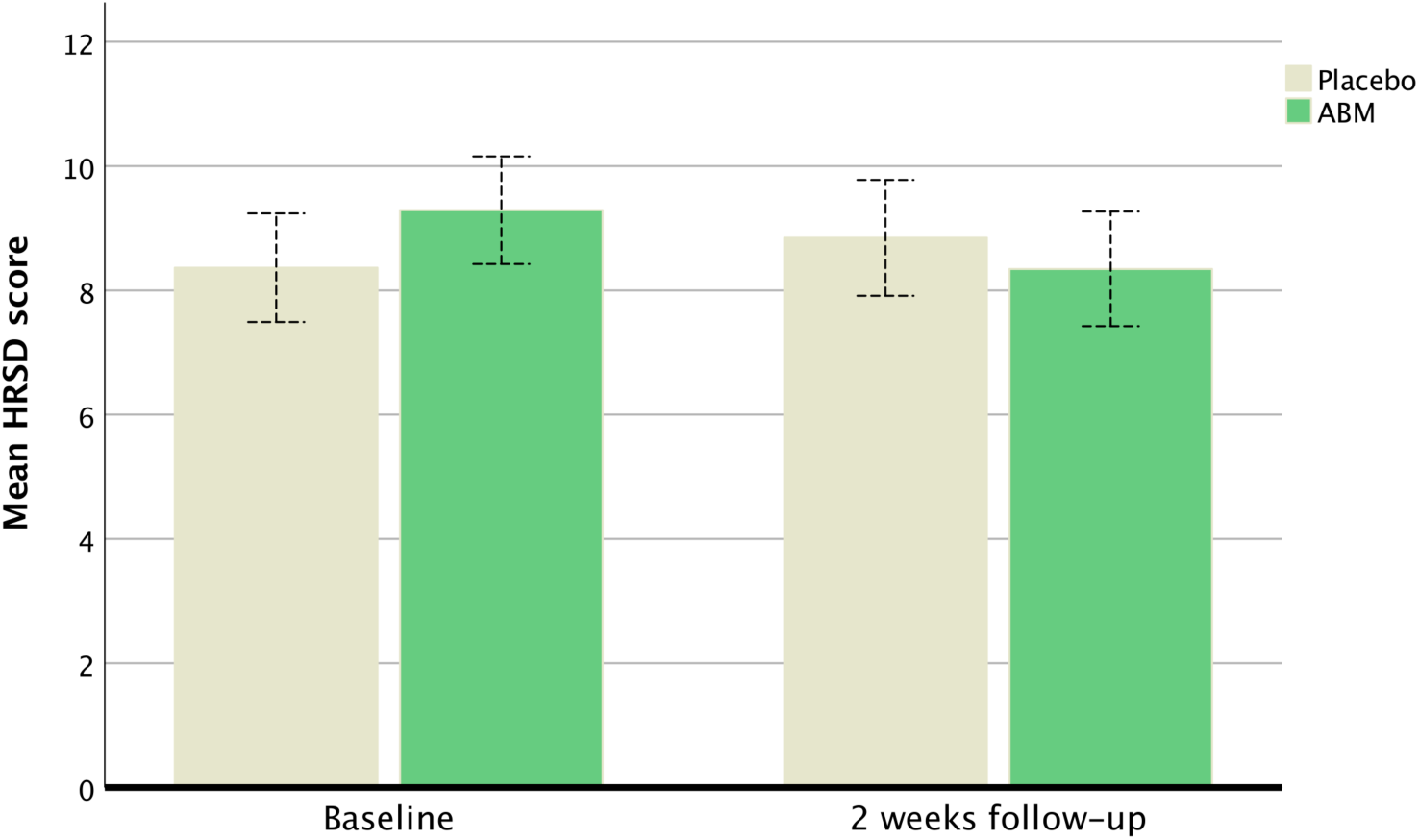
The mean symptom change in at baseline and after two weeks of ABM training for HRSD. Whiskers represent the 95% CI.

Although not statistically significant, there was a mean HRSD difference at baseline (Table 1.). A post hoc sensitivity test was therefore conduced by excluding potential outliers in the high end of HRSD scores (HRSD > 17 = cut-off for moderate depression). The post hoc test showed statistically significant differences in HRSD between ABM (n=135) and placebo (n=139). Means and standard deviations at baseline were 7.5 (3.9) for ABM and 7.2 (3.9) for placebo and changed to 7.3 (5.1) and 8.1 (5.4) at two weeks follow-up [F (1,272) = 4.48, η^2^= .02, p = .03] indicating that the observed changes in clinician-rated symptoms was not confounded by random variation at baseline.

**Table 1:** Demographic and sample characteristics per protocol. MDE=Major Depressive Episodes according to M.I.N.I. SSRI= any current usage of an antidepressant belonging to the Selective Serotonin Reuptake Inhibitors. ISCED= International Standard Classification of Education. P-values from Pearson Chi-Square test are presented for dichotomous variables.

|  | Placebo (n=148) | ABM (n=153) | p-value |
| --- | --- | --- | --- |
| Age | 41.5 (13.6) | 40.2 (12.7) | .68 |
| Gender (females) | 103 | 109 | .68 |
| Education Level (ISCED) | 5.9 (1.2) | 6.0 (1.1) | .79 |
| Medication (SSRI) | 43 | 38 | .43 |
| Number of previous MDE | 4.1 (4.6) | 4.1 (4.9) | .92 |
| Symptoms at baseline: |  |  |  |
| BDI-II | 13.6 (9.8) | 15.0 (10.6) | .23 |
| HRSD | 8.3 (5.1) | 9.2 (6.0) | .12 |
| BAI-II | 9.0 (7.4) | 9.6 (9.4) | .54 |

There was no statistically significant difference in self-reported depression as measured by the BDI-II between the ABM and the control condition [F (1,309) = 2.07, p = .15]. Both ABM and placebo showed statistically significant (time) improvement based on the BDI-II [F (1,309) = 67.77, η^2^= .18, p < .001]. Means and standard deviations at baseline were 14.9 (10.5) for ABM and 13.8 (9.7) for placebo and were 11.5 (10.4) and 11.4 (9.8) at two weeks follow-up (Figure 2.).

**Figure 2.**
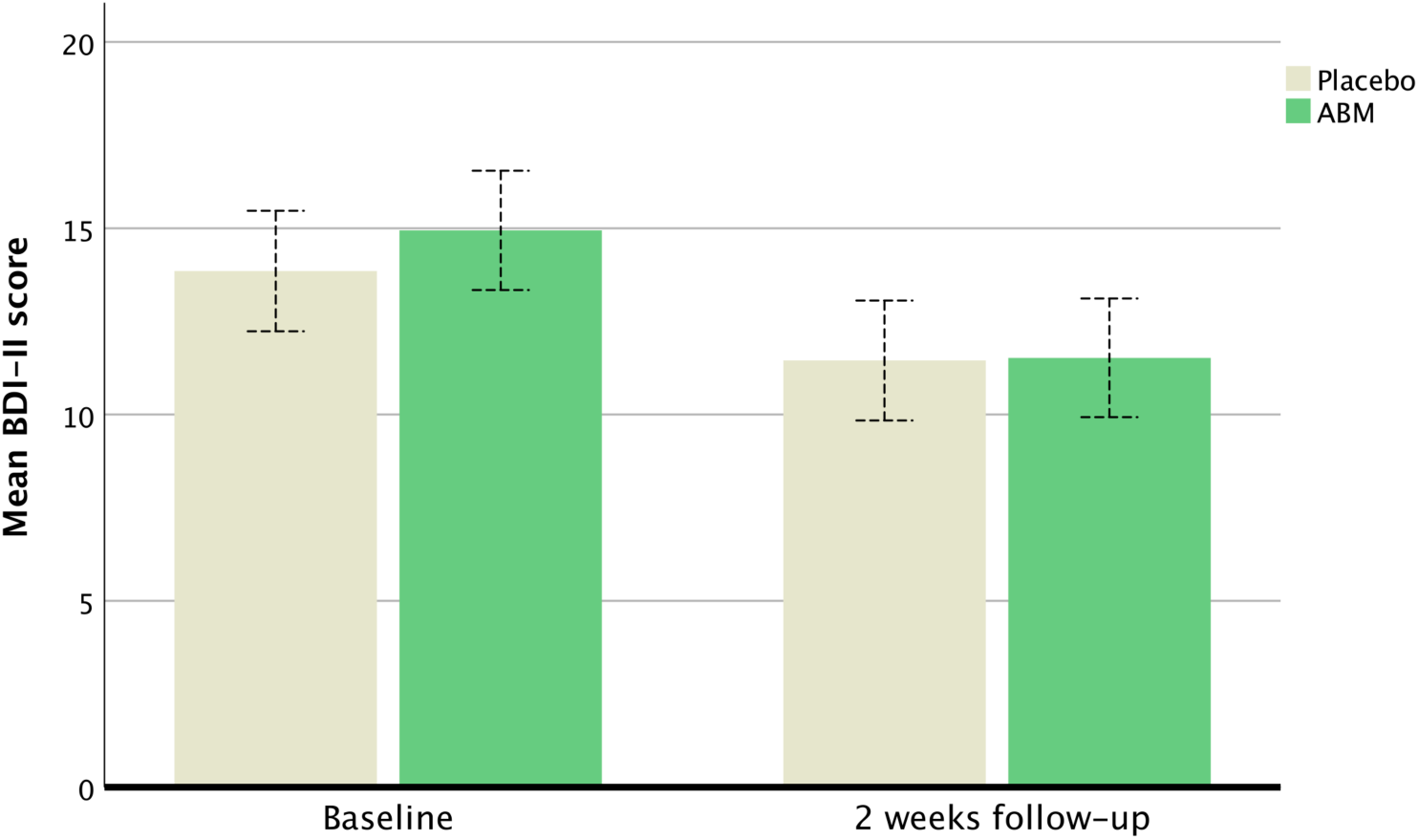
The mean symptom change in at baseline and after two weeks of ABM training for BDI-II. Whiskers represent the 95% CI.

### Symptom change after ABM and attentional biases

A combined factor repeated measure ANOVA revealed a statistically significant interaction between changes in AB (time) and changes in HRSD (time) meaning that an association between relatively more positive biases was associated with symptom change in both groups [F (1,309) = 3.793, η^2^= .01, p = .05]. A post hoc linear regression model showed that there was a statistically significant association between changes HRSD and changes and AB within the ABM group (Beta= .22, p < .00, R^2 = .05) but not within the placebo group (Beta= .06, p = .48) (Figure 3.).

**Figure 3.**
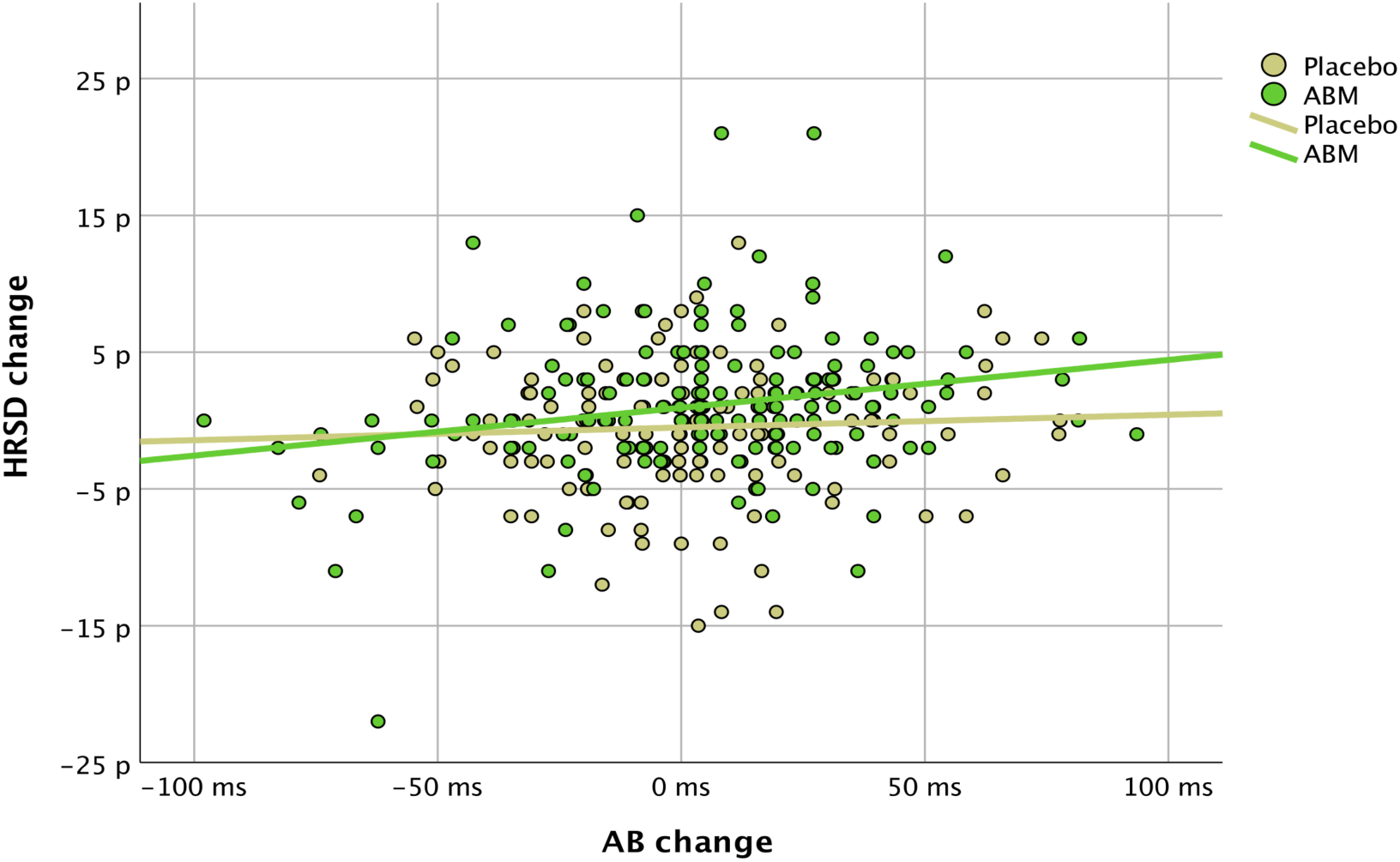
Relationship between changes in HRSD (two weeks follow-up minus baseline) and changes in AB (two weeks follow-up minus baseline). Positive values = changes towards more positive biases.

### Symptom specificity and depression status

No statistically significant differences (time) were found for self-reported anxiety as measured by the BAI-II [F (1,309) = 1.648, p = .20]. As found for self-reported depression (BDI-II) general symptom improvement was found in both ABM and in the placebo group [F (1,309) = 27.783, η^2^= .08, p < .01].

A repeated measure ANOVA with depression status (current MDD) and intervention as fixed factors revealed a statistically significant interaction between HRSD (time) and depression status [F (1,309) = 6.222, η^2^= .02, p = .01]. Means and standard deviations at baseline were 16.5 (6.1) for current MDD and 7.7 (4.5) for no current MDD and changed to 12.7 (7.3) and 7.6 (5.5) at two weeks follow-up. The interaction between HRSD and the intervention remained statistically significant [F (1,309) = 9.313, η^2^= .03, p < .01]. The three-way interaction between depression status, intervention and HRSD was not statistically significant [F (1,309) = 3.657, p = .06]. The results show that blinded rated symptoms was higher in current depression and decrease relative to placebo after ABM independent (although at trend level) of depression status.

## DISCUSSION

### Main findings

Two weeks of ABM training significantly reduced blinded clinician-rated symptoms as compared to a control condition in a group of patients previously treated for depression and with various degrees of residual symptoms. Relatively more positive change in AB was associated with symptom improvement. Corroborating cognitive models of emotional disorders, the degree of symptom improvement increased with degree of relatively more positive bias within the ABM group. A larger intervention independent effect was also found for self-reported depression and self-reported anxiety. Symptom improvement was dot driven by, but was more pronounced in participants who also fulfilled the formal diagnostic criteria for current depression. Post hoc exclusion of participants with high HRSD scores had no impact on the ABM effect, and per protocol analyses showed that the results are robust against imputations and compliance rates.

In addition to being among the strongest predictors for recurrence in depressive disorder (4) residual symptoms also cause significant functional impairments, manifested in a variety of domains, including work and leisure activities (18). The latter is important because one aim of depression treatment is to restore the patients’ previous level of functioning. However, residual symptoms in depression have traditionally not been the target of treatment trials. Given the recurrent nature of the illness, we suggest that this should change and that our study provides an example of how this may be done.

It is not clear why ABM versus placebo did not differ as measured by the BDI-II self-report scale, but instead showed a relatively large improvement in both the ABM- and placebo. In another recent study, fifty-two participants in a major depressive episode (MDE) was recruited for ABM training and a similar significant change in BDI-II in both the ABM- and placebo was found (19). The lack of association between ABM and self-reported symptoms was not expected from the pre-registered hypothesis and interpreting the source of this lack of evidence is speculative. However, self-report may be more influenced by placebo or expectation effects. A general effect of the intervention may also be linked to cognitive training effects present in both training conditions. An assessment only group will help in distinguishing placebo- from ABM in future studies. Symptom assessment based on both syndrome diagnoses, self-rated and blinded clinician-rated symptoms represent strength in the current study and the results clearly underpin the importance of comprehensive clinical evaluations in future research.

Participants completed the training in their homes, which is an advantage as it makes it more feasible as a treatment. Furthermore, compliance was high (see supplemental results) which is important and contrary to the conclusion in a recent review of meta-analyses (11). These authors conclude that ABM paradigms are most effective when delivered in the laboratory rather than at home. Specifying the scheduling of the training sessions individually might have increased motivation to do and focus on the task.

### Implications

The results suggest that ABM does indeed exert an effect on blinded clinician-rated symptoms. The finding would be considered clinically non significant in treatment trials. However, it is not clear how to interpret clinical relevance of small HRSD score changes in this group. The HRSD and BDI-II is only moderately correlated and may partly reflect different depression constructs (20). It is unclear whether the effect of ABM would increase and generalize across self-rated and clinician-rated symptoms if treatment continued for longer. Although residual symptoms are predictive of risk of relapse, it will be important to test in longer-term prevention studies whether the observed beneficial effect of ABM could translate into clinical relevance like reduction of relapse.

There is a pressing need to improve treatment and thus clinical trials should focus not only on efficacy, but also on identification of the underlying mechanisms through which treatments operate (21). We observed a number of people with large changes in AB that go along with symptoms improvement, but also with no change in HRSD or even worsening of HRSD scores. Identification of this individual variability may be useful in evaluations of treatment efficacy and in personalized treatment. The current study reports the pre-registered primary outcome, but stratification based on degree of AB, changes in dispersion and/or distribution of AB, or paradigms sensitive to the temporal expression of AB may help advance basic knowledge on the conditional benefit of ABM (22).

### Limitations

The study has several limitations that should be mentioned. Sparse research on this population impede calculations of sample sizes from prior studies but would further increase the stringency of the study design in accordance with CONSORT guidelines (23). Inclusion of 37 patients that also fulfilled the formal criteria for current depression represents a deviation from the pre-registered protocol. However, including current depression did not explain the results and increases the generalizability of the reported findings. Residual depression is widely defined as the study did not assess symptom change during former treatment and classify all symptoms as residual independent of treatment response.

### Conclusion

Previous studies have reported a mixture of positive and negative findings (14, 24-26) leading to a degree of scepticism about the impact of ABM on symptoms of depression (9). Our study is by far the largest randomized controlled clinical trial of ABM in a depressed population and gives a broader and more transparent picture of the true advantage of the intervention in this population. The results verify the proof of principle by showing that changes in AB is linked to changes in symptoms and suggest that ABM may have potential in the treatment of residual depression.

## DECLARATIONS

### Ethics approval and consent to participate

Written informed consent was obtained before enrollment. The study was approved by The Regional Ethical Committee for Medical and Health Research for Southern Norway (2014/217/REK Sør-Øst D). The authors assert that all procedures contributing to this work comply with the ethical standards of the relevant national and institutional committees on human experimentation and with the Helsinki Declaration of 1975, as revised in 2008.

### Consent for publication

Not applicable

### Availability of data and materials

The datasets used and/or analysed during the current study are available from the corresponding author on reasonable request.

### Authors’ contributions

NIL is PI in RCN, project number 247372 and in SEH, project number 2015052 and received funding for the projects. CJH is Co-PI in both projects. Both NIL and CJH were central in the study design, writing, and in revisions of the manuscript. VØH contributed in the planning of the project and the study design, and in revisions of the manuscript. EH, BK and LAM equally contributed in data collection, analysis, interpretation, writing and revisions. MB was central in data analysis, interpretations writing and revisions of the manuscript. TS and TB contributed in planning the study design, patient recruitment and diagnostics. RJ was post doctor and project manager for this study and was also responsible and directly involved in the data collection, recruitment, diagnostics, statistical analyses, interpretations, writing and revisions.

## Acknowledgements

We want to thank the following research assistants: Inger Marie Andreassen, Adrian Dahl Askelund, Dani Beck, Sandra Aakjær Bruun, Jenny Tveit Kojan, Nils Eivind Holth Landrø, Elise Solbu Kleven and Julie Wasmuth. Further, we thank Kari Agnes Myhre and Senior Consultant, Research leader; MD, Phd Erlend Bøen at Diakonhjemmet Hospital, Division of Psychiatry, for help and support during the recruiting period. We also thank our external recruitment sites, Unicare, Coperiosenteret AS, Torgny Syrstad, MD, Synergi Helse AS and Lovisenberg Hospital.

## Supplemental information

### Data reduction and compliance rates

Bias scores were calculated based on median reaction times. We choose to clean data for reaction times below 200- and 2000ms in accordance with recent literature that have aimed to grasp temporal stability in AB (27). The mean total trials included after applying the filter was 97.25% (,04%) in the placebo group and 97.23% (.02%) in the ABM group and did not differ between the groups [F (1,301) = .003, p = .958]. A second filter was used to exclude incorrect responses. The filter affected the placebo group by leaving 99.45% (.007%) and also 99.45% (.008%) in ABM group for further analyse and did not differ between ABM and placebo [F (1,301) = .002, p = .962]. A total of 13 participants lacked either pre- or post ABM data that form the basis for the calculation of bias scores. The series mean was used as imputation method for missing data.

Optimal ABM compliance was ensured using a calendar system describing the ABM training that was scheduled individually before the intervention. Compliance rates for the primary outcome measures per protocol (BDI-II and HRSD) were 100%.

Compliance rates (percentage of max 2688 trials) did not differ between ABM (M=83.5 (14.8)) and placebo (M=80.0 (21.2)) [F (1,300) = 2.731, p = .09]. A post hoc analysis restricted to participants with compliance above 50% (ABM n=147, placebo n=133) did not affect the association between HRSD change and ABM (M=85.4 (11.5)) versus placebo (M=86.4 (10.4)) [F (1,269) = 6.265, η^2^= .02, p = .01].

### Interaction between ABM and stimulus valence- and duration

There was a statistically significant interaction between HRSD change x face pairs (positive versus neutral, negative versus neutral, positive versus negative) x intervention (ABM versus placebo) that showed ABM more modification towards positive versus neutral stimuli as compared to the two other valence categories [F (2,309) = 3.278, p =.04]. Stimulus duration (500- or 1000 ms) did not interact with HRSD change and intervention [F (1,309) = .386, p = .53].

### Primary outcome per protocol

As in the ITT analyses, a repeated ANOVA showed a main effect of ABM as compared to placebo in HRSD scores when the 10 imputed ITT participants were excluded from the analysis [F (1,300) = 6.697, η^2^= .02, p = .01].

### Consort flow diagram

**Supplemental Figure 1.**
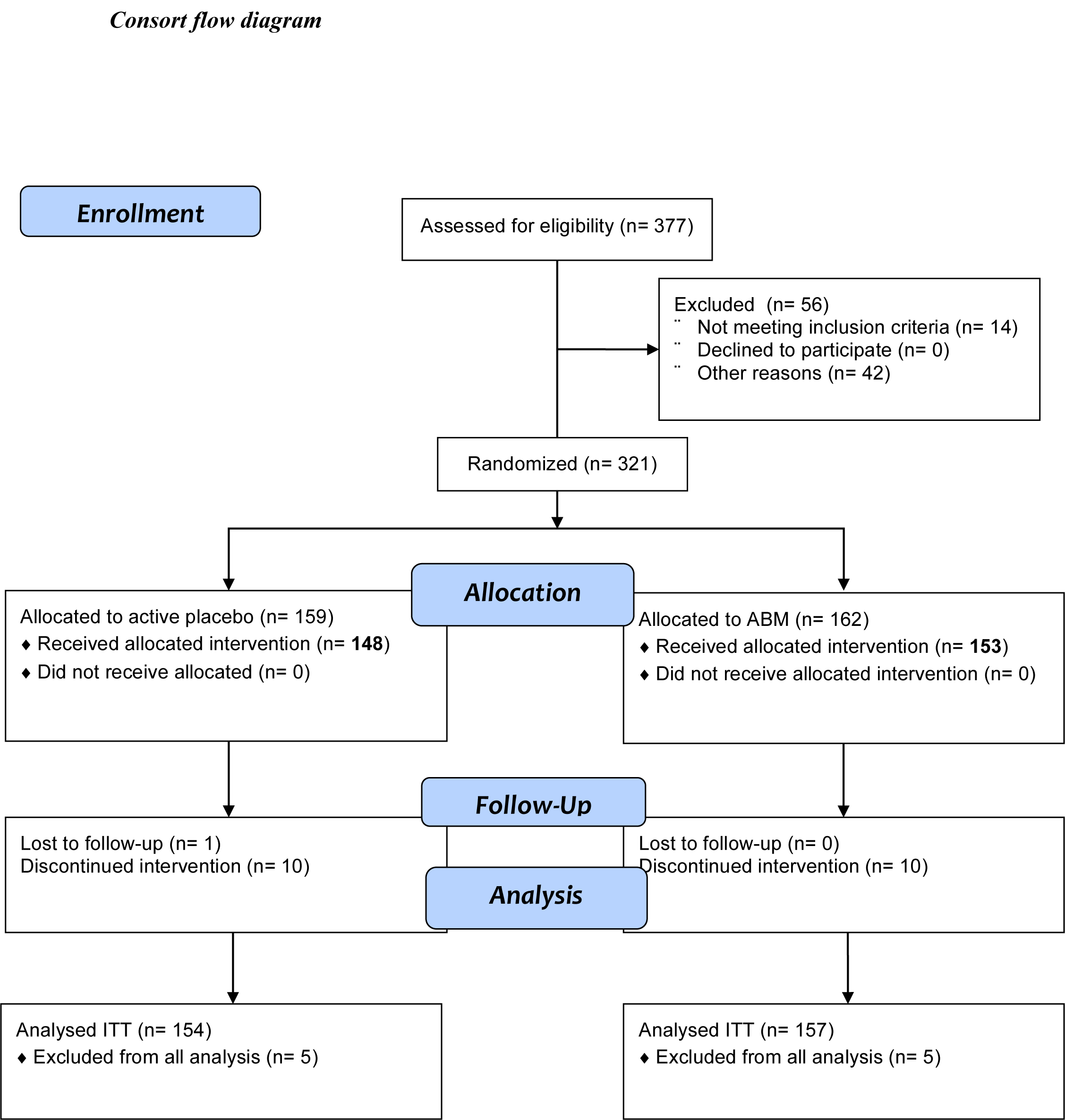
Flow diagram for enrolment, allocation to active placebo or ABM, follow-up after two weeks, and analyses in accordance with CONSORT (28). Not meeting inclusion criteria= no MDE according to M.I.N.I. Other reasons (n=42)= current or former mania and/or hypomania according to M.I.N.I. Ten participants (5 in each group) elected to have their data erased.

